# Cardiopulmonary bypass activates classical monocytes via shear-mediated activation of Store-Operated Calcium Entry

**DOI:** 10.1101/2022.05.04.490549

**Authors:** Weiming Li, Lan N. Tu, Lance Hsieh, Julian R. Smith, Yi-Ting Yeh, Anthony Sinyagin, Majid Ghassemian, Andrew Timms, Kevin Charette, David Mauchley, Michael McMullan, Lyubomyr Bohuta, Christina Greene, Mary C. Regier, Juan Carlos del Alamo, Ram Savan, Vishal Nigam

## Abstract

Patients undergoing cardiac surgery face significant inflammatory induced by exposure to cardiopulmonary bypass (CPB), contributing to heightened morbidity and mortality. The molecular and cellular mechanisms that underpin this inflammatory process remain unknown. To address this knowledge gap, we performed snRNA/ATAC-Sequencing on leukocytes from neonatal CPB patients. Classical monocytes become more prevalent and have dysregulation of inflammatory genes after CPB, indicating their role in CPB-associated inflammation. A genome-wide CRISPR screen and *in vitro* experiments in non-adherent monocytic cells identified two novel genes, SPTAN1 and RAF1, as effectors of hemodynamic stress. SPTAN1 and RAF1 activate store-operated calcium entry that results inflammation and cell death. snATAC-Seq revealed dynamically changing patterns of chromatin accessibility and AP-1 transcription factor binding after CPB exposure. These findings provide novel insights into the pathogenesis of CPB-associated inflammation, with broad implications for understanding the early stages of sterile inflammation and how non-adherent cells sense shear stress.

## Introduction

Over 30,000 pediatric^1^ and 150,000 adult patients^2^ undergo cardiac surgery every year according to the Society of Thoracic Surgeons Databases. Cardiopulmonary bypass (CPB) is routinely used during cardiac surgery to give the surgeon a bloodless field to operate in, while also minimizing ischemic damage to the body. In patients recovering from complicated cardiac surgeries, increased cytokine levels are associated with high mortality and extended intensive care stays^3^. Neonatal patients are particularly at risk, with a 10% mortality and a 30% complication rate^4, 5^. Despite CPB being used for over 70 years, there are number of open questions related to the pathogenesis of CPB associated inflammation that have slowed efforts to ameliorate this systemic inflammatory response. Understanding the underlying mechanisms by which CPB induces inflammation is critical for efforts to reduce post-cardiac surgery complications in patients.

CPB represents a unique opportunity to study how innate immune activation contributes to systemic inflammation since this process begins when cardiopulmonary bypass is initiated. Most of the clinical situations in which inflammation is typically studied — such as sepsis, lupus, rheumatic arthritis, and Kawsaki disease—involve studying the resolution phase, since the patients cannot be identified prior to the start of inflammation. This barrier represents significant challenges to understanding the early steps in inflammatory processes. In contrast, studying CPB patients could help answer important questions about the initiation of sterile inflammation.

CPB involves exposing the patients’ blood to a number of non-physiologic insults including supraphysiologic fluid shear stresses, which we have demonstrated are sufficient to induce cytokine expression and necroptosis in non-adherent leukocytes^6^. While previous studies examined how adherent cells^7^, red blood cells^8, 9^, and platelets^10^ respond to shear stress—it is unclear how non-adherent nucleated cells respond to shear stress. Understanding the mechanisms by which non-adherent cells respond to shear would address fundamental questions related to mechanobiology.

To identify the mechanisms that underlie CPB-associated inflammation, we performed single-cell sequencing of samples collected from neonatal CPB patients in conjunction with *in vitro* mechanistic experiments. We first performed snRNA/ATAC-Seq profiling of leukocytes from CPB patients, and identified classical monocytes as the largest population in the circulating PBMCs. An *in vitro* genome-wide CRISPR screen performed in human monocytic cells demonstrated that shear stress activates inflammatory responses via a mechanism that involves a non-erythroid spectrin (SPTAN1)/RAF1 signaling pathway. Additional mechanistic experiments demonstrated that SPTAN1/RAF1 promotes Ca^2+^ entry into the cell interior through the store-operated calcium entry (SOCE) pathway, consequently upregulating *Interleukin-8* (*IL8*), also known as *CXCL8*, expression. Patient snATAC-Seq provided a dynamic chromatin accessibility atlas during CPB-supported heart surgery, with over sixty percent of chromatin regions opened in response to CPB are located in promoter regions in PBMCs. In particular, we found several differential expression genes presenting increased or decreased accessibility at promoter regions in classical monocytes. Transcription factor (TF) motif enrichment analysis and *in vitro* experiments validated that the accessibility of binding sites of AP-1 TF members (JUN, JUNB, JUND, FOS, FOSL1 and FOSL2) is significantly promoted by CPB. Taken together, these data provides novel insights into the pathogenesis of CPB associated-inflammation and how non-adherent cells “sense” and respond to shear stress.

### CPB dramatic increases the proportion of classical monocytes in neonatal patients

To investigate how CPB modulates cellular composition and gene expression in an unbiased manner, we performed snRNA-Seq and snRNA/ATAC-Seq on PBMCs collected from neonatal patients (patient demographics in Supplemental Table 1) pre-CBP, at the end of CPB, 8 h, and 24 h post-CPB, (workflow outlined in Figure 1A). The pre-CPB samples served as a baseline control for each patient. 8 hours after CPB was selected as a time point since the majority of patients show signs of significant inflammation at this time. By 24 hours post-CPB, much of the inflammation has often resolved. We identified 13 distinct cell types from snRNA-Seq of 124,526 cells (Figure 1B), and 11 cell types from snATAC-Seq of 102,673 cells (Figure 1D), confirmed through the presence of marker genes and ATAC peaks (Figure S1-S2). Based upon quantification of cell counts for clusters derived from snRNA-Seq data, the percentage of classical monocytes increased by 28% at the end of CPB and by 116% 8h post-CPB as compared to pre-CPB, suggesting a pivotal role for classical monocytes in responding to CPB exposure (Figure 1C). Of note, other innate immunity-related cell clusters, CD8+ NKT-like and CD8+/CD4+ NKT-like cells, were also larger at the end of CPB and 8h post CPB as compared to pre-CPB. These results preliminarily indicated that the innate immune system was activated in patients exposed to CBP at the single cell level. Differentially expressed genes (DEGs) analyses of the snRNA-Seq data showed that CPB exposure broadly modulates the immune system at the level of gene expression regulation. We identified 48 to 405 DEGs in each cell type as compared to the pre-CPB time point (Figure 1E-G). Inflammatory and immune pathways were significantly modified in all of the cell clusters after CPB exposure (Figure 1H). Among them, classical monocytes displayed the strongest activation of positive regulation of immunity response and cytokine response genes based upon the number of genes and P-value. Notably, *ADAMTS3, EREG, CXCL8, MIR646HG*, and *PADI4* genes were initially identified as targets of CPB transcriptional regulation in total circulating leukocytes of CPB patients^6^. However, further investigation in this study revealed that these genes were specifically upregulated in monocytes (Figure S4). Hence, these findings underscore the pivotal involvement of classical monocytes in modulating the immune system response to CPB in neonatal patients.

**Figure 1.**
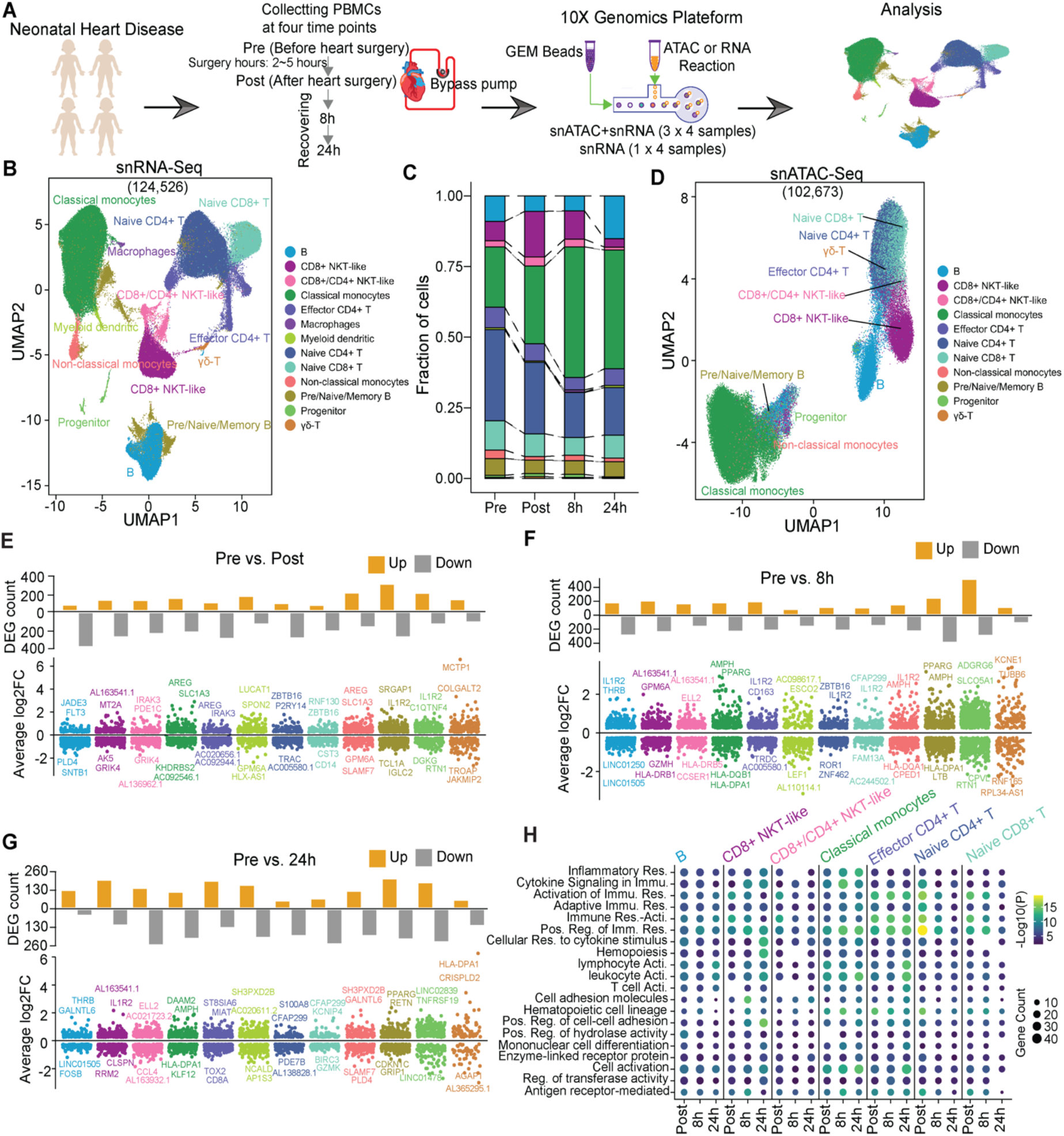
Profiling of snRNA-Seq and snATAC-Seq in neonatal cardiopulmonary bypass (CPB) patients. **A**. Schematic representation of PBMCs sampling from congenital neonatal CPB patients. PBMCs samples from four stages were collected in each patient. Pre represents before heart surgery; Post represents after heart surgery; 8h and 24h represent recovering stages. **B**. UMAP plot for cell clustering and cell type annotation of 124,526 cells based on snRNA-Seq. Thirteen cell types were annotated. **C**. Distribution of cell fractions of annotated cell types based on snRNA-Seq across four stages. **D**. UMAP plot for cell clustering of 102,673 cells based on snATAC-Seq and the cell type annotation mapping from snRNA-Seq. Eleven cell types were annotated. **E-G**. Differential expression genes (DEGs) were identified at Post, 8h, 24h time points, as compared to the baseline stage (Pre), respectively. The upper subpanel in each panel represents the count of DEGs. The bottom subpanel in each panel represents the distribution of average log2FoldChange of these DEGs in each cell type. **H**. Functional enrichment results of these DEGs detected in Post, 8h and 24h sampling points. Gradient yellow to blue color represent the value of -Log(P). The size of the dots represents the count of gene in each pathway. Statistical significance was accepted at P value less than 0.01. ‘Res’. means ‘response’; ‘Immu’ means ‘immune’; ‘Acti’ means ‘activity’; ‘Pos’ means ‘positive’; ‘Reg’ means ‘regulation’.

### Genome-wide CRISPR screen identified SPTAN1 and RAF1

Given that classical monocytes were the largest cluster of PBMCs after CPB exposure, we focused on monocytes for further mechanistic experiments. The expression of *IL8* was examined since we have previously shown that its expression increased in bulk RNA-Seq data from neonatal CPB patients^6^. Increased *IL8* levels have also been linked to worse outcomes for CPB patients^11, 12, 13, 14^. We found that *IL8* expression increased in classical monocytes at the end of CPB as compared to pre-CPB and subsequently decreased from this peak at the 8h and 24h post-CPB time points (Figure 2A). mRNA-Seq of RNA isolated from THP-1 cells, a human non-adherent monocytic cell line, exposed to *in vitro* shear stress in order to mimick CPB conditions also identified *IL8* as one of the genes significantly upregulated in response to shear stress (Figure 2B). Based upon these snRNA-Seq and *in vitro* data, we elected to use *IL8* expression as the readout for an unbiased genome-wide CRISPR loss-of-function screen to gain insight into how shear stress activates non-adherent monocytic cells.

**Figure 2.**
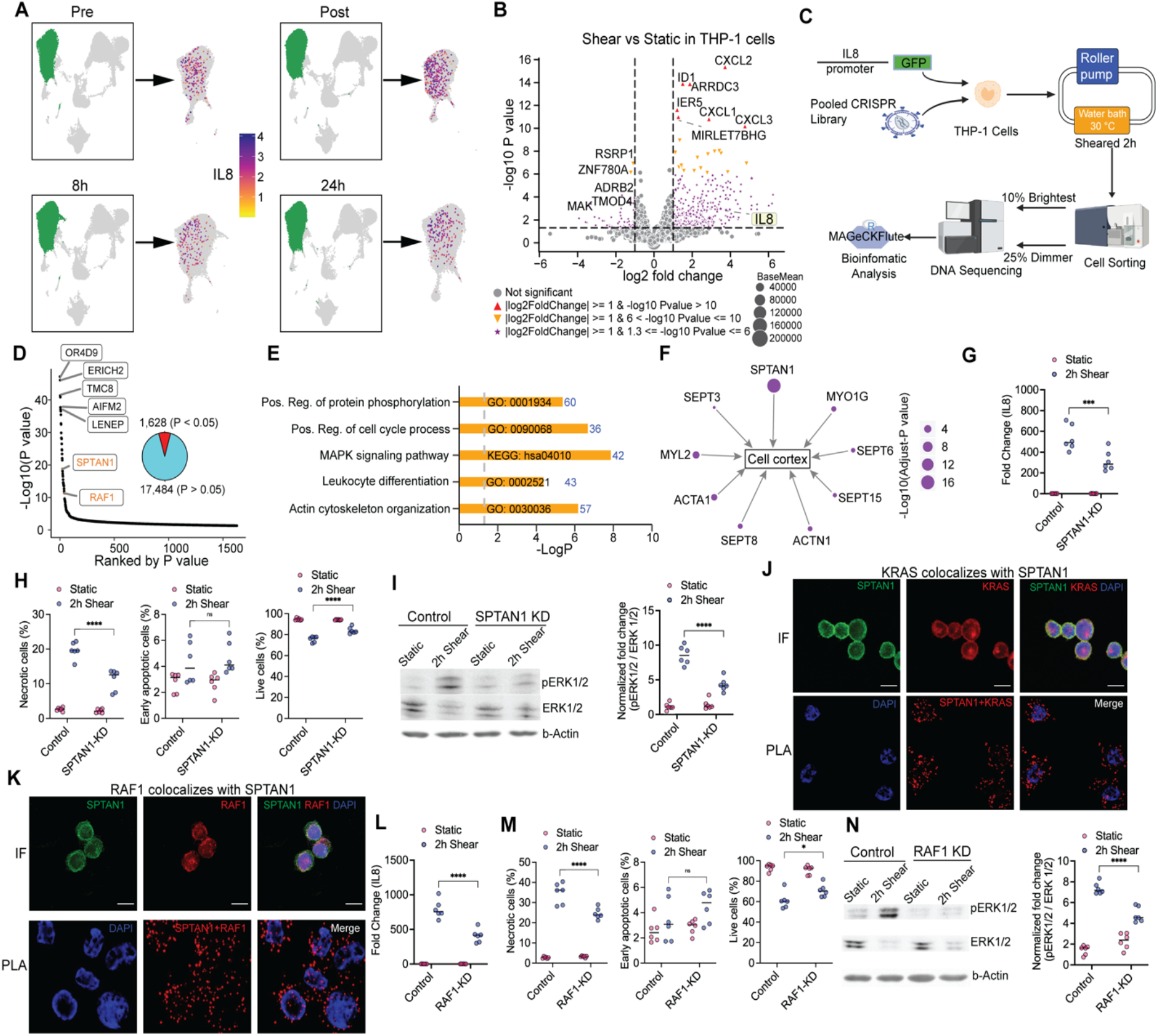
Genome-wide CRISPR screen identified two novel genes, SPTAN1 and RAF1, as candidates for the hemodynamic shear stress activation in non-adherent cells. **A**. UMAP plots of four stages highlighting classical monocytes with green color. Feature plots at right side of each UMAP plot displaying the *IL8* expression in four time points with blue dots repressing cells expressing *IL8*. **B**. Volcano plot for DEGs identified in sheared non-adherent monocytic cells (THP-1 cells) compared to static condition using mRNA sequencing. Three replicated were generated for each condition. Statistical significance was accepted with P value less than 0.05 and log2FoldChange less than -1or greater than 1. Seven top upregulated and five top downregulated genes were labelled. The important DEG IL8 was also labelled in a rectangle. Dots represent not significant. Red regular triangles represent DEGs with P value less than 1 x 10^−10^. Yellow inverted triangles represent DEGs with P value less than 1 x 10^−10^ but greater than 1 x 10^−6^. Stars represent DEGs with P value less than 1 x 10^−6^ but greater than 0.05. The size of dots represents the average of the normalized count values. **C**. Schematic representation of genome-wide CRISPR screen using the promoter of *IL8* as readout in THP-1 cells. **D**. Dot plot of CRISPR screen candidates ranked by P-value. Out of 1,628 genes, representing 8.5% of total, gRNAs were underrepresented in the 10% GFP bright cells with an adjusted P-value < 0.05. Top five genes and two novel important candidates were labelled. **E**. Selected key GO-terms from the Metascape analysis of 1,628 candidates. ‘Pos.’ means ‘positive’. ‘Reg.’ means ‘regulation’. **F**. Nine candidates associated with cortex pathway. The size of dot represents adjusted P-value. **G**. SPTAN1-KD THP-1 cells have 41.20% reduction in the shear-mediated activation of *IL8* expression as compared to sheared wildtype (WT) THP-1 cells based upon qPCR. Replicates N = 6. **H**. Quantification of cellular necrosis, early apoptosis, and live cells demonstrates a 43.40% reduction in cellular necrosis and 10.30% increase in live cells after shear in the SPTAN1-KD THP-1 cells as compared to control sheared cells. Replicates N = 6. **I**. SPTAN1-KD cells have decreased shear-mediated ERK1/2 phosphorylation as compared to control THP-1 cells. Replicates N = 3. ** P < 0.01, **** P < 0.001. **J**. KRAS colocalizes with SPTAN1 in THP-1 cells based upon immunocytofluorescence (IF) and proximity ligation assay (PLA). **K**. RAF1 colocalizes with SPTAN1 in THP-1 cells based upon immunocytofluorescence (IF) and proximity ligation assay (PLA). **L**. RAF1-KD cells have 48.50% reduction in shear-mediated *IL8* expression as compared to sheared control cells. Replicates N = 6. **M**. Quantification of cellular necrosis, early apoptosis, and live cells demonstrates that RAF1-KD cells undergo 29.30% less shear mediated necrosis and have 15.80% increased cell survival as compared to control THP-1 cells. Replicated N = 6. **N**. Knockdown RAF1 decreases the shear-mediated ERK1/2 phosphorylation as compared to control THP-1 cells. Replicated N = 3. * P < 0.05, ** P < 0.01.

The design of this CRISPR screen (outlined in Figure 2C) used the proximal *IL8* promoter driving green fluorescent protein expression (*IL8-GFP*) as a reporter in THP-1 cells. THP-1 cells were transduced with lentiviruses driving a genome-wide CRISPR/Cas9 interference library^15^ and then exposed to shear stress. The cells were sorted based on GFP brightness and then sequenced. Genes whose gRNAs were underrepresented in the brightest GFP pool were classified as candidate genes important for the shear stress response. We identified 1,628 candidate genes with an adjusted P < 0.05 as cutoff (Figure 2D and Supplemental Table 2). To our surprise, genes previously identified as shear stress sensors in endothelial cells—such as *PECAM1, VEGFR2 (KDR)*, and *PIEZO1* ^7, 16, 17^—were not identified as candidates in our CRISPR screen. The absence of endothelial “flow sensor genes” suggests that non-adherent circulating leukocytes utilize a different mechanism to sense shear stress as compared to adherent cells.

Given the number of candidate genes identified, we utilized gene-ontology term (GO-term) analyses to obtain a more nuanced understanding (Supplemental Table 3). The GO-term ‘Mitogen-activated protein (MAP) kinases (P = 2.30 x 10^−6^)’ was included in the result of functional enrichment analysis for these candidates. Several kinases in this subgroup—namely, MEKK2, MEK2, JNK3, and MEK— were previously identified as being critical to fluid stress activation of THP-1 cells^6^, which helped validate the results of the screen. The GO-term related to actin cytoskeleton was also one of the most significantly enriched (P = 2.30 x 10^−7^) (Figure 2E and F). This pathway contains nine CRISPR screen genes potentially associated with shear stress: *SPTAN1* (nonerythroid spectrin, P = 3.00×10^−17^), *MYL2* (myosin II light regulatory chain, P = 3.00 x 10^−4^), *ACTA1* (α-actin, P = 1.00 x 10^−3^), *MYO1G* (myosin 1g, P = 1.00 x 10^−2^), *Septin8* (P = 1.00 x 10^−2^), *ACTN1* (α-actinin, P = 2.00 x 10^−2^), *Septin6* (P = 2.00 x 10^−2^), *Septin15* (P = 3.00 x 10^−2^), and *Septin3* (P = 4.00 x 10^−2^) (Figure 2F). Specifically, the *SPTAN1* gene functions as an essential scaffold protein, stabilizing the plasma membrane and organizing intracellular organelles^18^, suggesting its potential importance in responding to shear stress.

To further study the role of *SPTAN1* in mediating the response of non-adherent THP-1 cells to fluid shear stress, we used CRISPR to generate knockdown *SPTAN1* (*SPTAN1-KD*) cells. *IL8* expression, cellular necrosis, and Extracellular signal-regulated kinase (ERK)1/2 phosphorylation were used as readouts since these markers have been shown to be activated by hemodynamic stresses^6^. qPCR showed a significant decrease in shear stress-induced *IL8* expression *in SPTAN1-KD* as compared to control THP-1 cells (Figure 2G). In addition, flow cytometry after Annexin V and Propidium iodide staining indicated reduced cellular necrosis in sheared *SPTAN1-KD* THP-1 cells as compared to control cells (Figure 2H and Supplemental Figure S6A). Western blot analysis confirmed reduced ERK1/2 phosphorylation in *SPTAN1* deficient cells (Figure 2I). Together, these results demonstrate that *SPTAN1* plays a key role in mediating shear stress responses of non-adherent cells, which, to our knowledge, represents a non-canonical function for the *SPTAN1* gene.

### RAF1 helps mediate SPTAN1’s shear stress activation of non-adherent cells

While previous studies have shown that *SPTAN1* can activate ERK signaling via a SRC-dependent mechanism^19^, we did not identify SRC as a candidate in our CRISPR screen (Adjusted P = 0.80). In addition, inhibition of SRC signaling did not blunt the shear activation of *IL8* expression (Supplemental figure S5A) in THP-1 cells. These data suggest an alternative pathway is involved in *SPTAN1* modulation of ERK activity. *RAF1*, a member of the MAPK pathway, was also identified as significant in the CRISPR screen (Adjusted P = 7.88 x 10^−15^) (Figure 4I). The MAPK signaling pathways, including the RAS-RAF-MEK-ERK signaling cascades, regulate fundamental and diverse cell processes^20^. RAF1 activates MAP kinase-kinase (MEK), which in turn phosphorylates and activates mitogen-activated protein kinase (MAPK), also known as extracellular signal-regulated kinase (ERK)^21, 22^. This signaling pathway prompted us to examine whether KRAS/RAF1 signaling is a mechanism by which *SPTAN1* stimulates ERK activity in the context of hemodynamic shear stress. KRAS was selected as the RAS family member to be studied since it had a P-value of 0.06 in the CRISPR screen. We observed that both KRAS and RAF1 co-localized with SPTAN1 based upon immunocytofluorescence (IF) and proximity ligation assays (PLA) (Figure 2J-3K). *RAF1-knockdown* (*RAF1-KD*) cells were found to have decreased flow-mediated *IL8* expression (Figure 2L) and necrosis (Figure 2M and Supplemenatal Figure S6B) as compared to wild-type cells. Fluid shear stress-mediated phosphorylation of ERK was also decreased in RAF1 deficient cells, as determined by immunoblot (Figure 2N). Taken together, these data suggest a novel interaction between KRAS/RAF1 and SPTAN1 that plays an important role in shear stress-mediated cytokine expression and necrosis.

### Shear stress facilitates Ca^2+^ entry through a SPTAN1/RAF1 mechanism

We sought to elucidate the mechanisms through which hemodynamic shear stress modulates calcium signaling, since extracellular calcium is necessary for fluid stress activation of non-adherent cells^6^. Additionally, our analysis of DEGs from snRNA-seq revealed enrichment for calcium-responsive genes towards the end of CPB (**Figure S5**). Using phosphoproteomics, we identified proteins that were phosphorylated in response to shear stress (Supplemental Table 4). Stromal Interacting Molecule (STIM) family members (STIM 1 and 2) were phosphorylated in response to shear. Specifically, STIM1 phosphorylation between amino acids 616-634 increased by over 80% after 30 minutes of shear stress exposure (P < 7.00 x 10^−60^). STIM2, identified as a significant gene in our CRISPR screen (P = 5.98 x 10^−6^), presented an increase in phosphorylation between amino acids 717-730 of over 23% after 30 minutes of fluid stress exposure (P < 5.24 x 10^−58^). STIM1 and 2 play a key role in store-operated calcium entry (SOCE) since they migrate to the inner surface of the cell membrane where they form a complex with calcium release-activated calcium channels (ORAI) to facilitate calcium entry. The region in which we found increased STIM1 phosphorylation has been shown to stimulate SOCE via a ERK1/2 depenent mechanism^23^, thereby promoting calcium influx.

In order to demonstrate that shear stress activates SOCE, we utilized a bimolecular fluorescence complementation (BiFC) STIM1/ORAI1 Venus reporter^24^ in which Venus signal develops when tagged STIM1 and ORAI1 are in close proximity. Sheared STIM1/ORAI1 Venus THP-1 cells have increased Venus’s signal (10.6% +/- 0.7 Venus positive) as compared to static control (1.8% +/- 0.2) (Figure 3A and B). In addition, knocking down either *STIM1* or *ORAI1* significantly reduced responsiveness to hemodynamic stress compared to control cells, as measured by both IL8 expression and cell necrosis (Figure 3C-D and Supplemental Figure S6C). Together, these data suggest that STIM1 and ORAI1 play an important role in the hemodynamic activation of inflammatory genes in response to shear stress.

**Figure 3.**
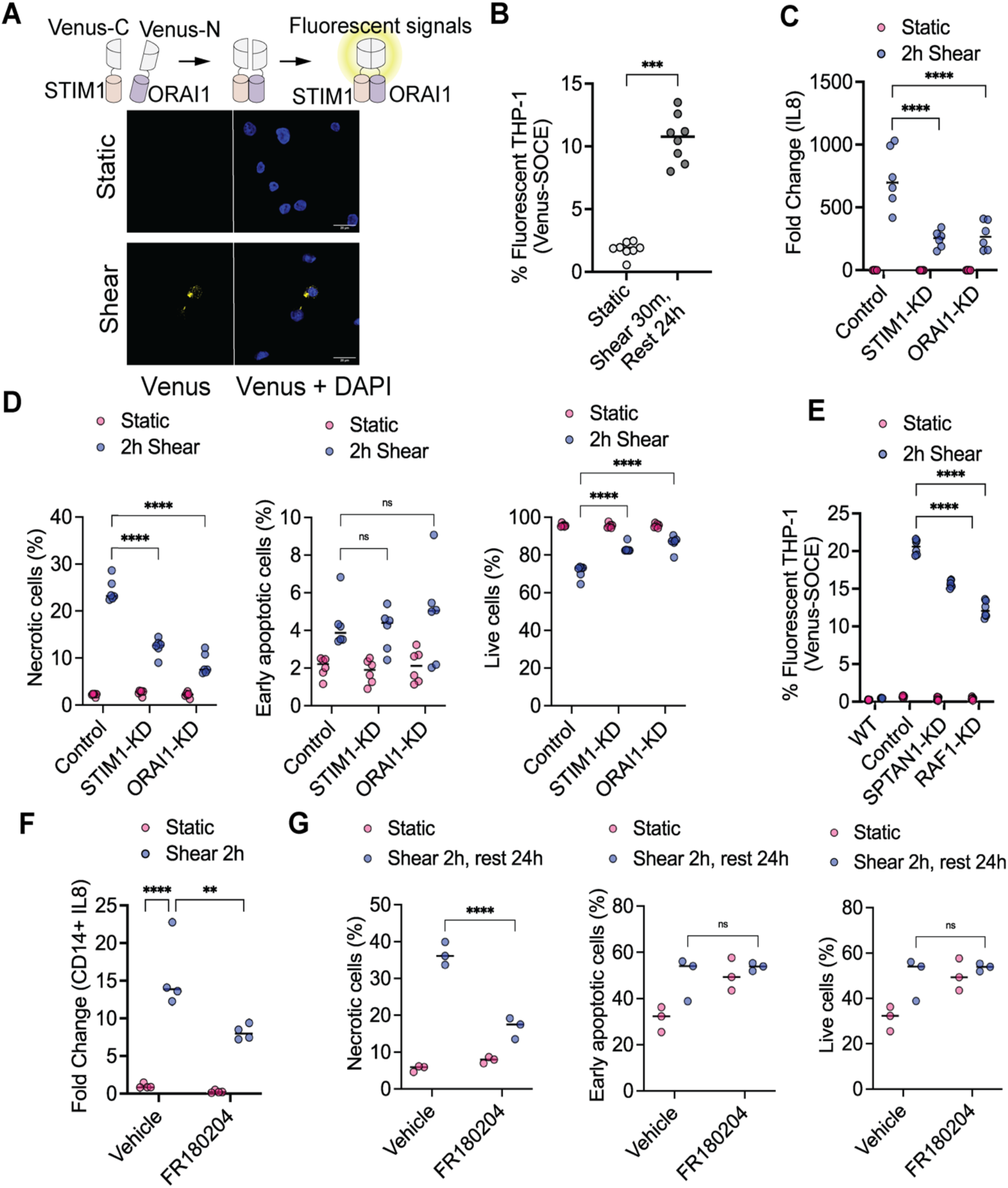
Hemodynamic shear stress promotes calcium entry through activating Ca2+ entry pathway (SOCE) via SPTAN1/RAF1 signaling. **A**. Visualization of the interaction of STIM1 and ORAI1 using split Venus as yellow signal reporter in bimolecular fluorescence complementation (BiFC) assay. A yellow signal indicates STIM1/ORAI1 interaction after shear stress compared to the static condition in THP-1 cells. **B**. Quantification of the Venus positive BiFC STIM1/ORAI1 THP-1 cells between sheared and static conditions. Shear stress condition possesses 10.60% +/- 0.70% Venus positive cells while static condition only having 1.80% +/- 0.20% Venus positive cells. **C**. STIM1-KD and ORAI1-KD cells have decreased shear mediated *IL8* expression as compared to control THP-1 cells. Replicates N = 6. **D**. Quantification of flow cytometry data showing STIM1-KD and ORAI1-KD cells have decreased shear mediated cellular necrosis and have increased cell survival. Replicates N = 6. **E**. SPTAN1-KD and RAF1-KD cells have decreased STIM1 and ORAI1 interaction based on decreased split Venus STIM1 and ORAI1 signal. Replicates N = 6. **F**. FR180204, an ERK inhibitor, decreased shear stress mediated *IL8* expression in CD14+ primary monocytes. **G**. Quantification of flow cytometry data displaying a decrease in shear stress mediated necrotic cells following treatment with FR180204 in CD14+ primary monocytes.

Considering that ERK1/2-mediated phosphorylation of STIM1 is crucial for the store-operated calcium entry (SOCE) pathway^23^, and that our data indicates that SPTAN1 and RAF1 regulate ERK1/2 signaling, we next examined if SPTAN1 and RAF1 are involved in the shear activation SOCE by using the STIM1/ORAI1 Venus cells. *SPTAN1* and *RAF1-KD* cells displayed decreased shear mediated Venus signal, demonstrating that SPTAN1 and RAF1 are upstream of SOCE in the context of fluid stress (24% reduction and 40.3% reduction, respectively) (Figure 3E). These results suggest a close linkage of hemodynamic stress activation of STIM1/ORAI1 with SPTAN1 and RAF1, identified by CRISPR screen. Furthermore, our experiments indicate the significance of SOCE in regulating both flow-mediated gene expression and cell death.

In order to examine the effect that shear stress and shear-mediated ERK signaling have on primary monocytes, we treated CD14+ primary human monocytes with FR180204, an ERK inhibitor, prior to exposing the cells to shear stress. Shear stress exposure is sufficient to upregulate *IL8* expression (Figure 3F) and activate cellular necrosis (Figure 3G and Supplemental Figure S6D) in vehicle treated CD14+ cells. Furthermore, shear stress induced *IL8* expression and cellular necrosis were significantly reduced in the sheared, FR180204 treated CD14+ primary human monocytes compared to vehicle control (Figure 3F and G). These results demonstrate a similar regulatory gene pattern between THP-1 cells and CD14+ primary human monocytes, providing evidence that our findings are not limited to immortalized cell lines.

### CPB conditions alter chromatin accessibility and promote JUN binding

Changes in chromatin accessibility can lead to alterations in gene expression patterns, affecting various cellular and molecular processes. We hypothesized that chromatin accessibility can be dynamically regulated to modulate gene expression and cellular responses in responding to shear stress in classical monocytes. To understand how exposure to CPB regulates gene expression via altering chromatin accessibility, we conducted snATAC-Seq and bulk ATAC-Seq experiments to assess the impact of CPB on chromatin accessibility. snATAC-Seq data from the patients shows dynamic opening and closing of ATAC peaks in response to CPB (Figure 4A). Over 60% of the differentially modulated ATAC peaks are in the promoter regions of genes (Figure 4B). These data indicated that the shear stress experienced during CPB is capable of inducing alterations in chromatin accessibility, thereby modulating gene expression. Given the critical role of classical monocytes and the high enrichment of differential peaks at promoters, we focused our examination on promoter peaks (Figure 4C). In classical monocytes, we identified twelve genes exhibiting increased chromatin accessibility at their promoter regions, accompanied by upregulated gene expression following CPB (Figure 4D and Figure S8C). In contrast, we found thirteen genes with decreased chromatin accessibility at their promoter regions, accompanied by downregulated gene expression following CPB (Figure 4E and Figure S8D). To further test this premise, we performed bulk ATAC-Seq on sheared THP-1 cells and static controls. For open peaks, 212 peaks (38.20%) of 555 snATAC peaks in patients were supported by bulk ATAC-Seq in sheared THP-1 cells (Figure 4F). For closed peaks, 304 peaks (23.62%) of 1,287 peaks in patients were confirmed in shear THP-1 cells (Figure 4F). For example, we found that ATAC peaks of *DUSP8* and *TREML4* at promoter region opened at the end of CPB and with shear while the peaks *MS4A4A* closed under both conditions (Figure 4G). To understand which transcription factors interact with the differentially modulated ATAC peaks, we conducted motif enrichment analyses on the snATAC-Seq data from classical monocytes. AP-1 transcription factors—including JUN, FOS, and BATF—opened following CPB and closed at 24h after CPB (Figure 4H). A similar enrichment for AP-1 factor binding was seen in the sheared THP-1 cells (Figure 4I). Since previously we had found that shear stress induces nuclear translocation of JUN transcription factor in a calcium dependent manner^6^, we performed transcription factor footprinting analyses on the patient and THP-1 ATAC-Seq datasets. The results demonstrated that the signals of transcription factor binding activity of the JUN and FOS TF families were enhanced in both patient snATAC-Seq and THP-1 bulk ATAC-Seq data (Figure 4J-5K and Figure S10). Additionally, CUT&RUN experiments using a JUN antibody detected 125 open and 62 closed peaks in response to shear stress in THP-1 cells (Figure S9), further emphasizing JUN as a sensitive transcription factor sensing fluid shear stress. These data suggest that CPB alters chromatin accessibility that contributes to gene regulation.

**Figure 4.**
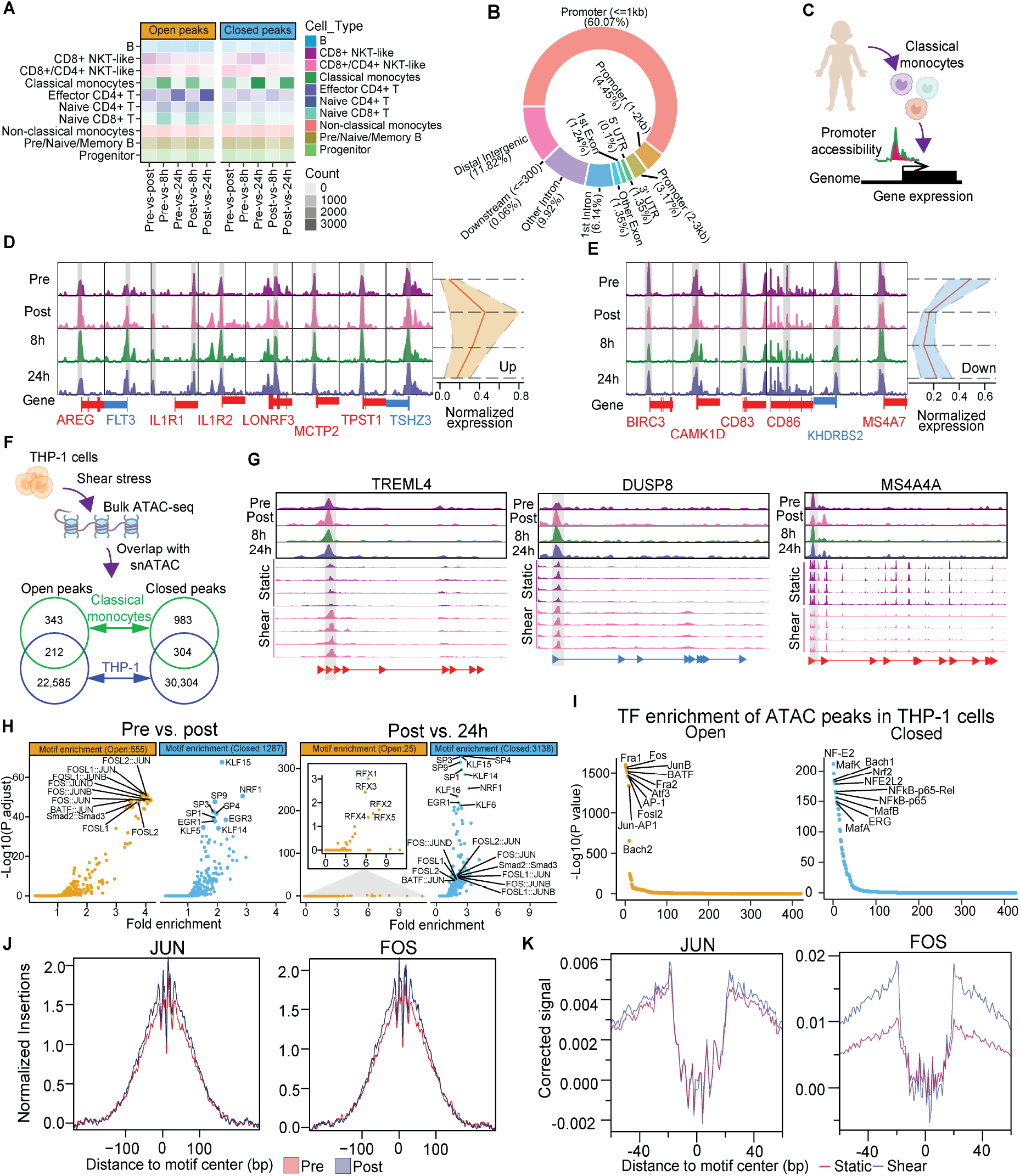
Shear stress induced shifts in chromatin accessibility at promoter regions and mediated transcription factors (TFs) binding events in classical monocytes. **A**. Heatmap presented the count of open and closed peaks identified in Post, 8h, and 24h time points across all cell types in PBMCs from CPB patients. **B**. Structure annotation of these peaks detected in all cell types in PBMCs from CPB patients. Over 60.07% peaks were located on promoter region. **C**. Schematic representation of how shear stress modulates promoter accessibility influencing gene expression in classical monocytes of CPB patients. **D**. Promoters accessibility of eight genes were opened, accompanying upregulated gene expression after CPB. Subsequently, both chromatin accessibility and gene expression tend to recover at 8h and 24h time points. **E**. Promoters accessibility of six genes were closed, accompanying downregulated gene expression after CPB. Subsequently, both chromatin accessibility and gene expression tend to recover at 8h and 24h time points. **F**. Schematic representation of bulk ATAC-Seq in THP-1 cells. Venn plots are showing the overlapped open and closed peaks between classical monocytes of CPB patients and THP-1 cells. **G**. Three candidate genes, *TREML4, DUSP8*, and *MS4A4A*, exhibit chromatin accessibility shifting at the promoter region in classical monocytes, a finding supported by THP-1 cells. **H**. AP-1 TFs, including JUN, JUNB, JUND, FOS, FOSL1 and FOSL2, were significantly enriched in open peaks at Post time point while significantly enriched in closed peaks at 24h time point. Top TFs were also labelled. **I**. TF enrichment analysis of open and closed peaks detected in THP-1 cells. Top 10 TFs were labelled in open and closed plots, respectively. **J**. Footprinting analysis of JUN and FOS TFs in classical monocytes. The signal of JUN and FOS increased at Post time point in CPB patients. **K**. Footprinting analysis of JUN and FOS TFs in THP-1 cells. Same pattern of these two TFs were observed in THP-1 cells.

## Discussion

In this paper, we have used several unbiased approaches to address open questions related to how CPB instigates systemic inflammation, especially the contributions of the shear stress present in the CPB circuit. snRNA-Seq data from neonatal CPB patients identified how CPB exposure modulates PBMC populations and the gene expression profiles within each cluster. Through a genome-wide CRISPR screen and phosphoproteomics, we have identified a novel pathway by which shear stress activates Ca^2+^ signaling in non-adherent cells. snATAC-Seq and *in vitro* ATAC-Seq data demonstrate that CPB conditions are sufficient to alter chromatin accessibility. These data may lead to focused inventions to reduce CPB associated inflammation and other conditions in which leukocytes are exposed to supraphysiologic shear stresses, such as aortic or carotid stenosis. In addition, our findings will advance the understanding of fundamental questions related to how cells “sense” shear stress.

The single cell sequencing data presented here provides the first comprehensive view of how PBMC populations change in response to CPB exposure. Exposure to CPB modulated differential gene expression across all the cell populations (Figure 1E-G) and the DEGs in each cluster correlate to terms related to inflammation and activation of immune cells (Figure 1H). We focused on the classical monocyte cluster for deeper examination given that it was the largest cluster at the post-CPB time points (Figure 1C). While we have found significant shifts in the populations of PBMCs after CPB exposure, further work is necessary to elucidate the mechanism that underpin these changes and the contributions of specific leukocyte populations. Two potential mechanisms driving a higher percentage of classical monocytes after CPB exposure are migration of monocytes into the peripheral circulation or the loss of other cell populations, due to cell death and/or migration out of the circulation. Emergency myelopoiesis, which involves migration of monocytes into the circulation in response to inflammatory responses, is a possible mechanism for the shift in the CPB-mediated leukocyte populations that needs be studied in future work. Studies are ongoing aimed at elucidating the mechanisms by which myeloid cells are mobilized and determining what the functional phenotypes of the mobilized cells are in terms of inflammatory modulation. In addition, some of the changes in gene expression and chromatin accessiblity observed in classical monocytes from patients but not reproduced in the *in vitro* shear experiments could be the result of monocytes mobilizing out of the bone marrow, with gene expression and chromatin patterns related to monocyte maturation. If circulating myeloid cells that enter circulation after CPB exposure propagate inflammation, efforts to limit the shifts in the circulating leukocyte populations would be beneficial.

There is growing interest in how biomechanical stimuli modulates leukocytes^25^. We and other groups have found that shear stress can activate Ca^2+^ signaling in myeloid cells^6, 26, 27^. In contrast to previous candidate-based approaches ^26, 27^ which studied the contributions of PIEZO 1 and 2 to the shear stress response, we propose here a novel shear stress responsive pathway consisting of SPTAN1/RAF1/SOCE/AP-1 signaling that is based on unbiased screens and confirmatory experiments. Our CRISPR interference screen identified SPTAN1 and RAF1 as significant contributors in the shear activation of *IL8*. Immunofluorescence and proximity ligation data suggested that SPTAN1 colocalizes with KRAS and RAF1, and we propose that this interaction is a key part of the shear stress responsive pathway. CRISPR based knockdown of *SPTAN1* and *RAF1* resulted in decreased shear activation of *IL8* expression, cellular necrosis, and ERK1/2 phosphorylation. Potentially, shear causes conformational changes in spectrin that promote KRAS/RAF1 interaction, thereby activating RAF1 and downstream ERK1/2 signaling. This premise is based upon data that shear stress has been shown to cause conformational changes of red blood cell spectrin^9, 28^ and that SPTAN1 can be a scaffold for protein kinases, such as SRC^19^. ERK1/2 mediated phosphorylation of STIM1^23^ promotes its translocation to the cell membrane where it interacts with ORAI1 resulting in Ca^2+^ entry via SOCE. Our phosphoproteomic and BiFC data implicated STIM1/ORAI1 as having a role in the shear response. We have recently demonstrated that JUN, an AP-1 transcription factor, translocates into the nucleus after shear stress via a Ca^2+^ dependent mechanism^6^. So, the Ca^2+^ influx could induce AP-1 transcription factor binding to chromatin as demonstrated by the ATAC motif, DNA footprinting, and CUT&RUN data. The transcription factor binding may result in the shear stress responsive gene expression. To our knowledge, our data is the first to suggest that SPTAN1 and RAF1 interact after shear stress contributing to activate the pathway of calcium entry. More broadly, having the cell cortex GO-term being a hit in the CRISPR screen leads to the premise that shear stress imposes loading on the cell cortex, which in turn activates downstream kinases that activate SOCE.

The data presented in this paper will assist efforts to develop targeted treatments to limit CPB-associated inflammation. Currently, pediatric CPB patients are routinely administered preoperative steroids to limit inflammation. However, a recent large randomized clinical trial demonstrated that prophylactic use of steroids did not improve outcomes^29^. Our data presents several therapeutic targets that have not been previously studied in the context of CPB. Specifically, RAF1, ERK1/2, and SOCE can be targeted with drugs. Sorafenib, for example, is an FDA approved drug that can inhibit RAF1^30, 31^. There are ongoing clinical trials using BVD-523 (Ulixertinib), which is believed to be a selective ERK1/2 inhibitor, in cancer patients^32^. FDA approved drugs have also been identified as inhibitors of SOCE^33^. Further research is necessary to identify drugs that could ameliorate CBP-associated inflammation by targeting the SPTAN1/RAF1/ERK/SOCE signaling pathway that we have implicated in this paper.

More broadly, the mechanistic experiments in this paper provide novel insight into how cells “sense” shear stress. In contrast to the extensive research looking at how shear modulates adherent cells like vascular endothelial cells, there is limited understanding of how non-adherent nucleated cells “sense” shear stress. In addition to the genes that we have validated in this paper and in previous work^6^, it is worth noting that our CRISPR screen also identified *CAV1, PECAM1*, and *VEGFR3*. These genes have been shown to be involved in modulating the shear response of endothelial cells^7, 34^. Although a number of candidate genes identified in our CRISPR screen have been validated, nearly 1,600 candidate CRISPR screen genes and could be fertile ground to gain deeper insight into the mechanisms that underlie the shear stress response. Our whole genome CRISPR screen is one of the first efforts to use an unbiased method to identify genes critical for shear stress in PBMCs, with the sole other published effort looking at endothelial cells^35^. The shear responsive pathway identified in this paper also has implications for other clinical situations in which blood is exposed to increased shear stress, such as aortic valve and carotid artery stenoses, along with other situations in which mechanical support of the circulation is required, e.g. ECMO and LVAD. In conclusion, we have used single cell profiling of neonatal PBMCs in conjunction with unbiased *in vitro* experiments to address important questions regarding the pathogenesis of CPB-associated inflammation along with how cells “sense” and respond to shear.

## Acknowledgements

We thank Dr. Eric Evans and Dr. Tricity Andrew for advice and revise on our manuscript. We would also acknowledge the University of Washington Institute for Stem Cell and Regenerative Medicine Core.

## Author contribution

V.N. conceptualized and supervised this project. W.M.L. performed snRNA/snATAC, bulk RNA-Seq, bulk ATAC-Seq and CUT&RUN analysis. Y. T. Y performed CUT&RUN experiments. L.S., L.N.T, and J.S. performed the experiments. M.G. performed the mass spec experiments and related bioinformatics. A.T. performed the bioinformatic for the CRISPR screen. K.C., D.M., M.M., L.B., and C.G. helped provide the patient samples. M.R. performed the snRNA/ATAC-Seq. R.S. supervised J.S. and helped edit the manuscript. V. N. and W.M.L. organized and wrote the manuscript.

## Data and materials availability

All of the snRNA, snATAC, bulk RNA-Seq, CRISPR screen, bulk ATAC-Seq and CUT&RUN datasets have been deposited in Gene Expression Omnibus under accession numbers GSE262146, GSE213184, and GSE213184. The remaining data are available in the main text or the supplementary materials.

## Highlights

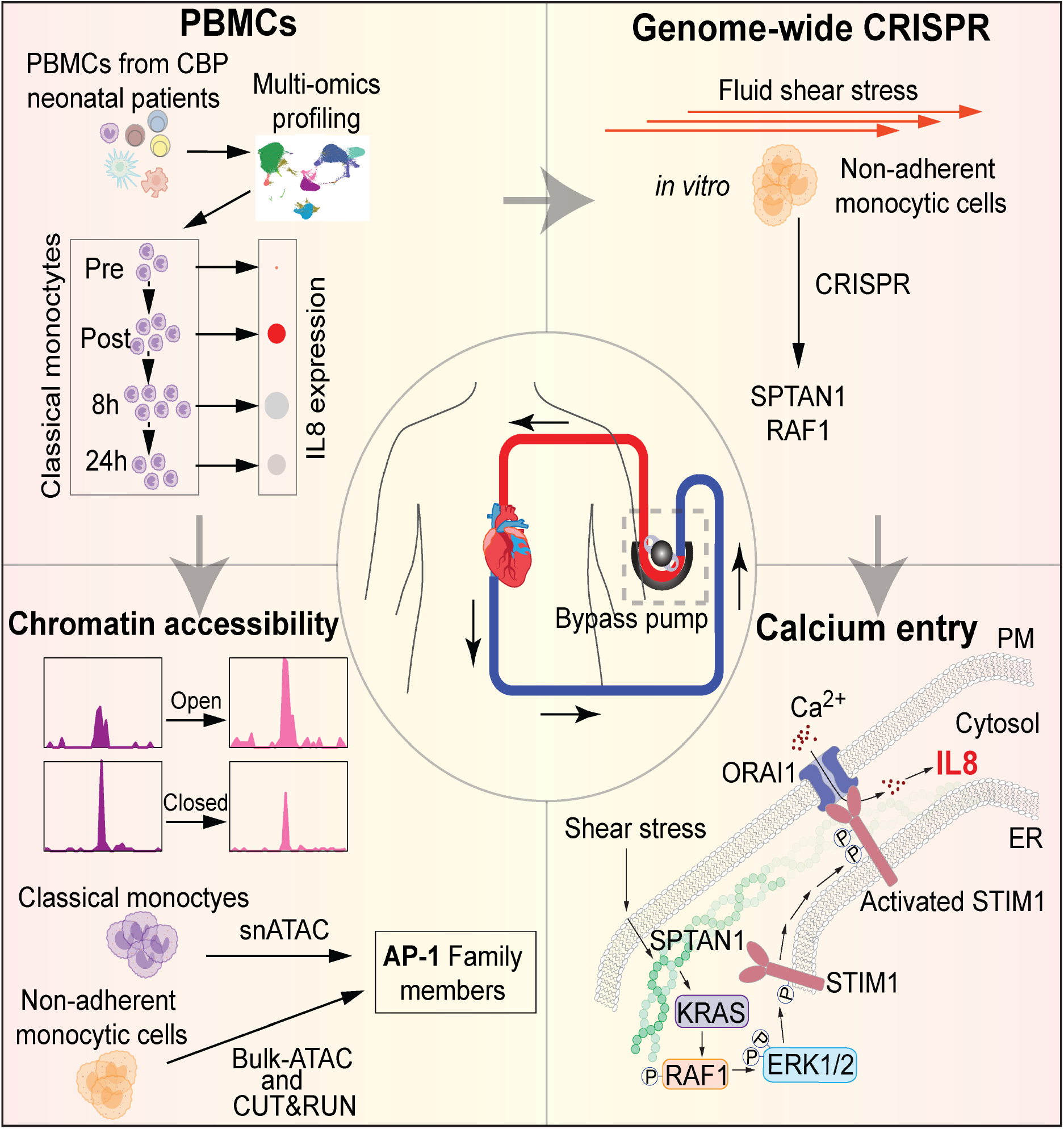

- The supraphysiologic shear stresses of CPB activate inflammatory programs in circulating classic monocytes
- The shear stress activates SPTAN1/RAF1 signaling in non-adherent monocytic cells
- Increased phosphorating STIM1 facilitates Ca^2+^ entry through SOCE pathway in non-adherent monocytic cells
- AP-1 transcription factors binding chromatin to drive pro-inflammatory gene expression and necroptosis

