## supplemental text and figures for "Cardiopulmonary bypass activates classical monocytes via shear-mediated activation of Store-Operated Calcium Entry"

**The PDF file includes:**

Figures. S1 to S10
Materials and Methods
Tables S1 to S4
References

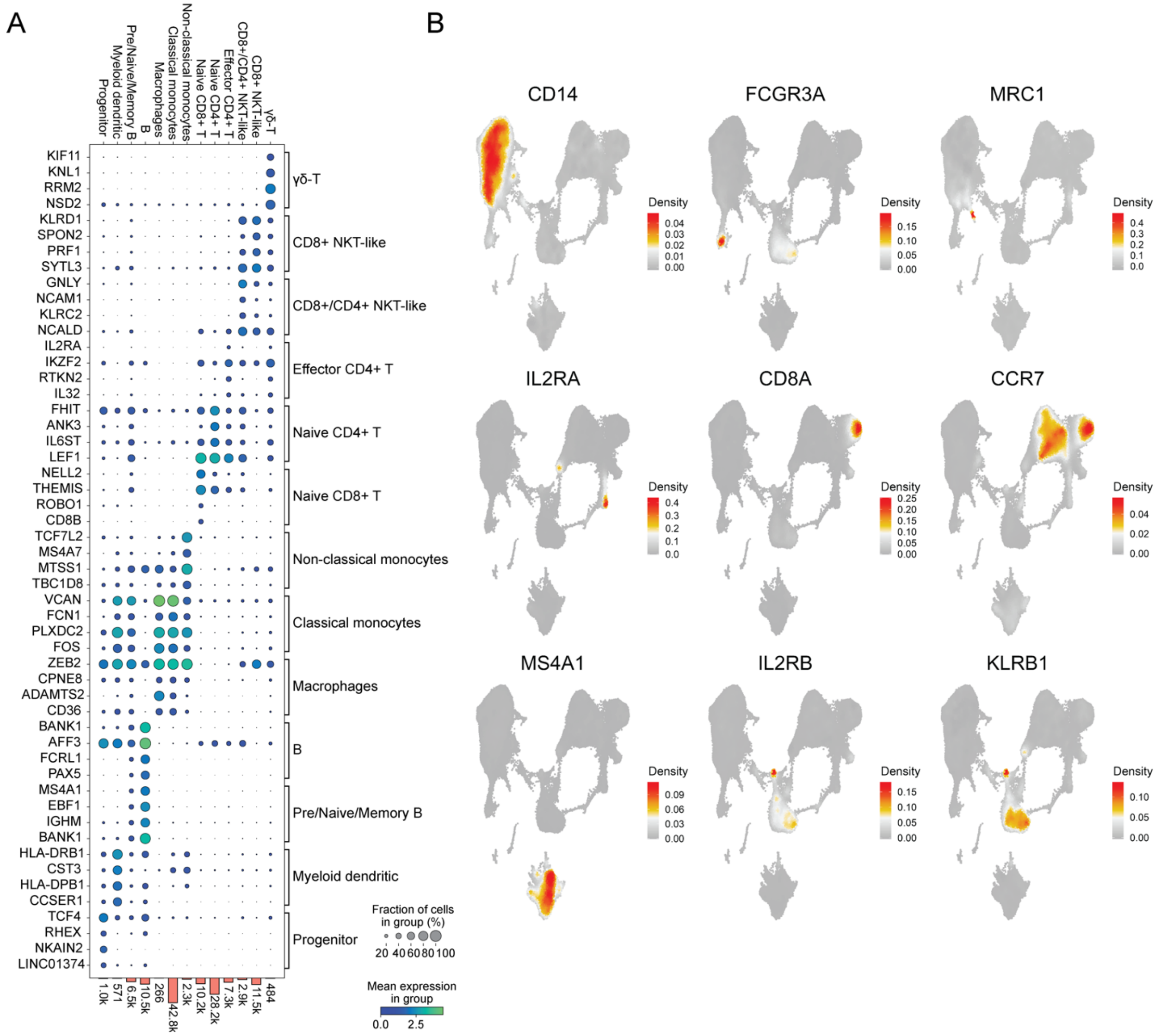


**Figure S1. The markers of snRNA were used to identify the cell types in PBMCs. A.** Dot plots were showing the relative gene expression of top four markers in each cell type. The bar graphs represented the count of cells in each cell type at bottom. The size of dot represented expressed fraction of cells in group. Gradient blue color depicted the mean expression of marker in group **B.** Density feature plot of gene expression displaying nine specific markers in PBMCs.

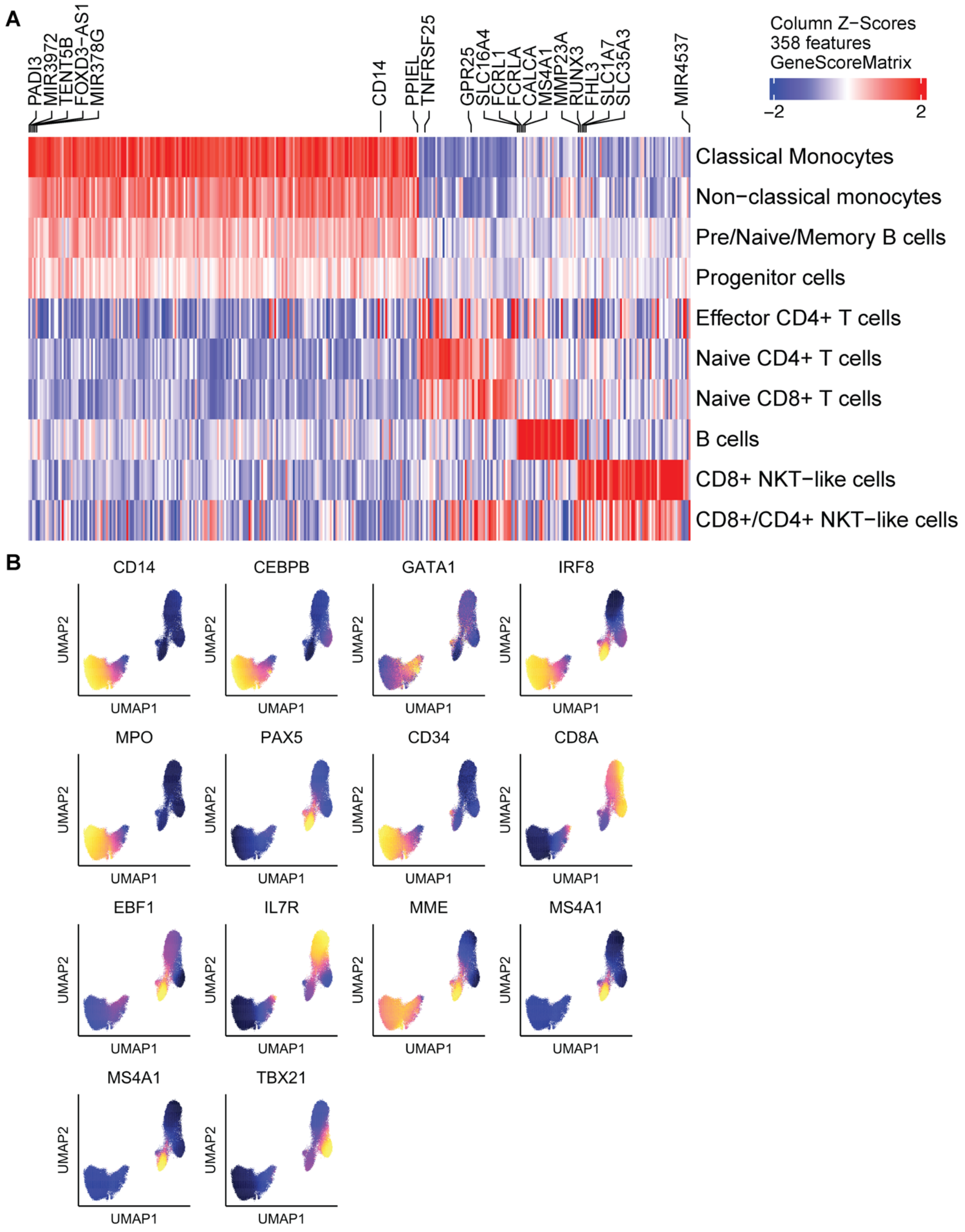


**Figure S2. Cell types were identified using snATAC. A.** Heatmap was showing gene scores of 358 features across all cell types. **B.** Gene scores of fourteen specific markers confirming cell types in snATAC were exhibited in feature plots.

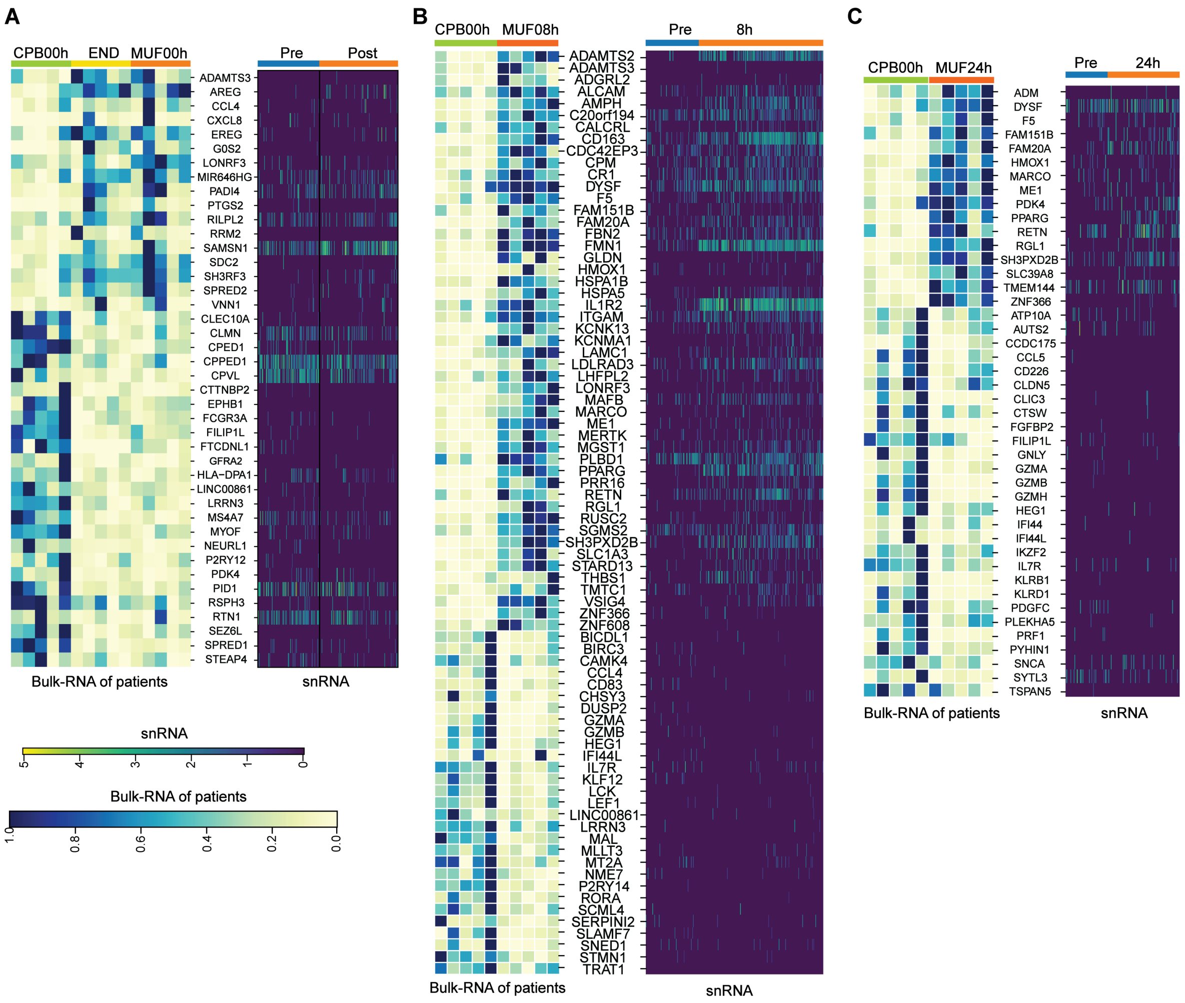


**Figure S3. Overlapped DEGs were identified between classical monocytes of snRNA and bulk-RNA patients (published at 2021). A.** Overlapped forty-two DEGs between the comparison of pre vs. post (right panel) from snRNA and the comparison of CPB00h vs. END/MUF 00h (left panel) from bulk-RNA of patients were presented in heatmap. **B.** Overlapped seventy-eight DEGs between the comparison of pre vs. 8h (right panel) from snRNA and the comparison of CPB00h vs. MUF 08h (left panel) from bulk-RNA of patients were shown. **C.** Overlapped forty-four DEGs between the comparison of pre vs.24h (right panel) from snRNA and the comparison of CPB00h vs. MUF 24h (left panel) from bulk-RNA of patients were displayed. CPB00h time point was identical to pre time point. END/MUF 00h was identical to post time point. MUF 24h was identical to 24h time point.

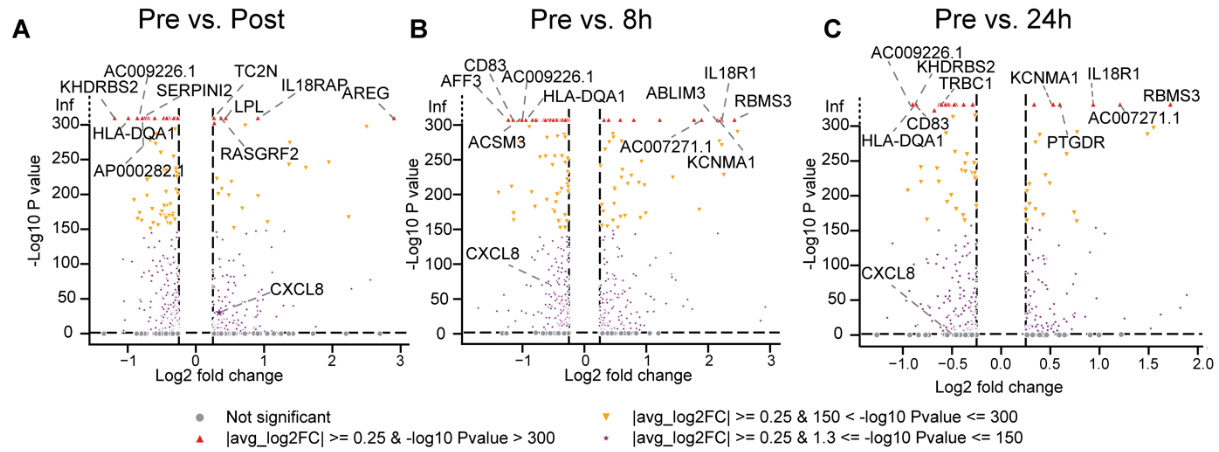


**Figure S4.** Volcano plots of differential expression genes (DEGs) from the snRNA-Seq dataset at post (panel **A**), 8h (panel **B**) and 24h (panel **C**) stages compared to pre stage in classical monocytes of snRNA. Top five upregulated and downregulated DEGs, as well as IL8(CXCL8) gene, were highlighted.

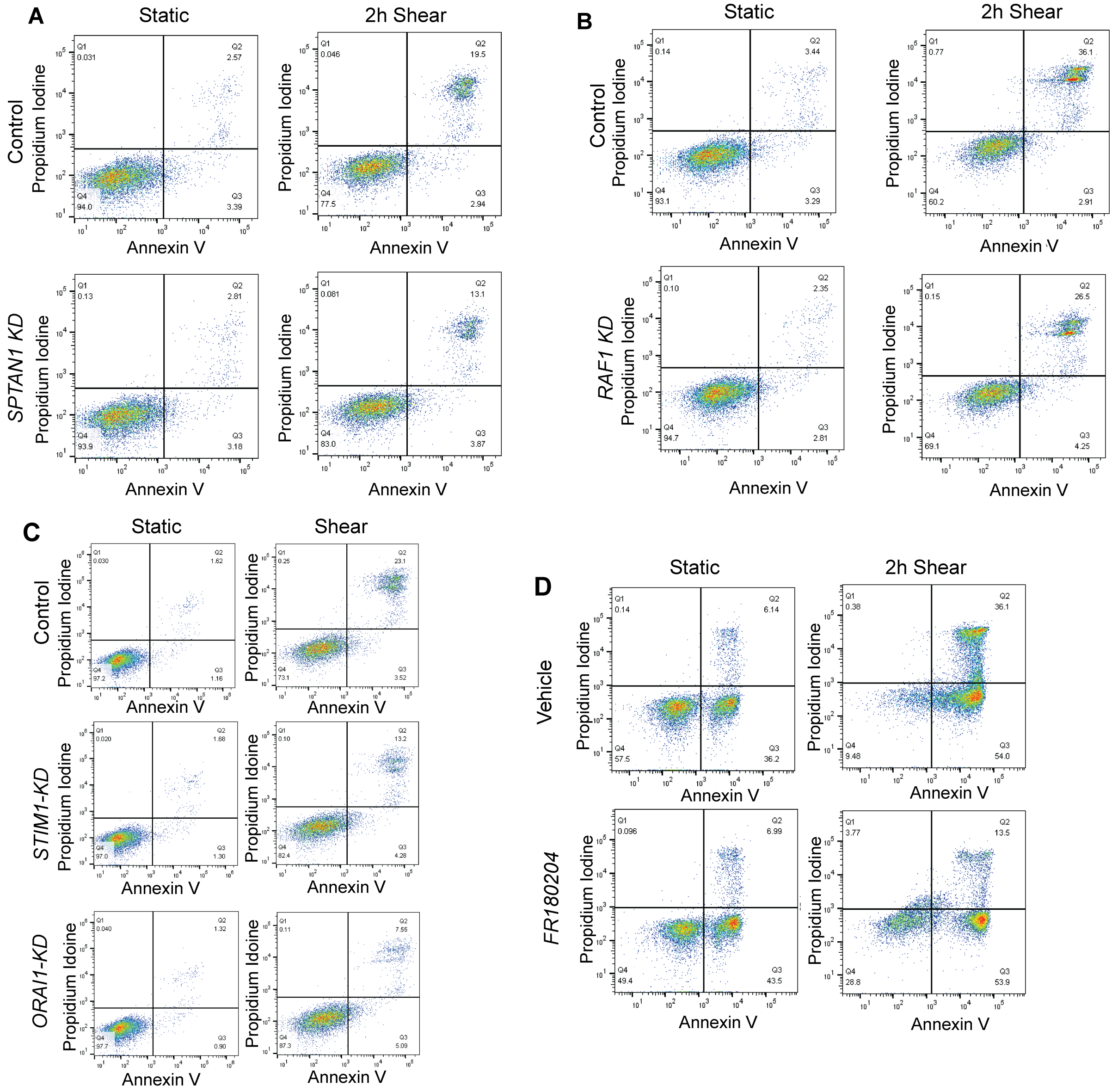


**Figure S5. Flow cytometry results of SPTAN1-KD, RAF1-KD, STIM1-KD, ORAI1-KD and FR180204 inhibitor conditions as compared to wild type (WT) or vehicle control in THP-1 cells. A.** Flow cytometry using PI and Annexin V staining demonstrated decreased shear mediated cellular necrosis (Q3) in SPTAN1-KD THP-1 cells as compared to WT control cells (N = 6). **B.** Flow cytometry demonstrated that RAF1-KD cells undergo 29.3% less shear mediated necrosis and have 15.8% increased cell survival as compared to control THP-1 cells (N = 6). **C.** Flow cytometry demonstrated that STIM1-KD and ORAI1-KD cells have decreased shear mediated cellular necrosis and have increased cell survival (N = 6). **D.** Flow cytometry demonstrated that ERK inhibited by FR180204 reduced shear stress induced IL8 expression and cellular necrosis in $CD14^{+}$ primary human monocytes (N = 6). KD means knockdown.

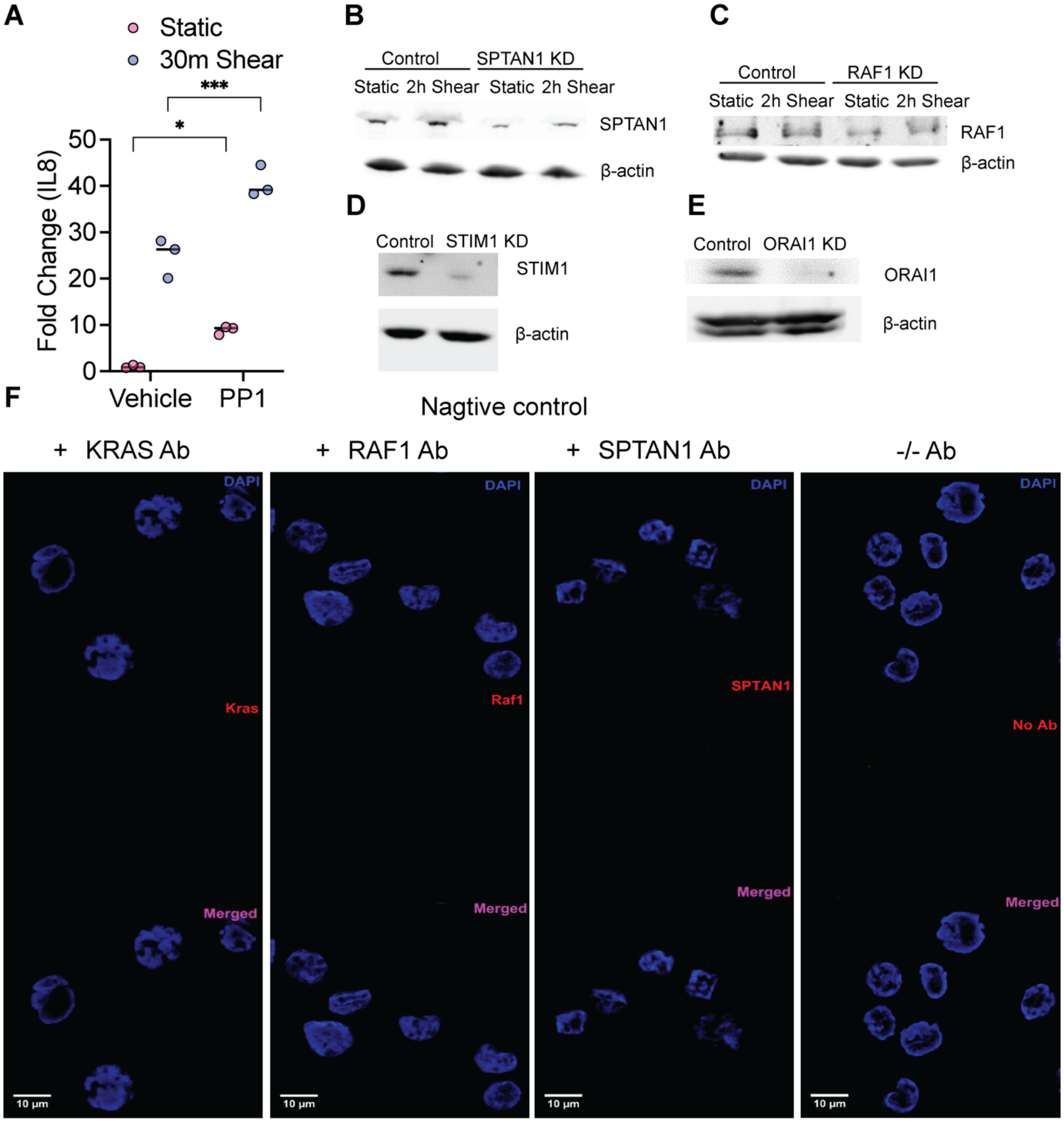


**Figure S6. A.** Src inhibitor PP1 does not significantly reduce the shear activation of *IL8* expression. THP-1 cells were treated either with 25 uM of PP1 or DMSO (Veh). The cells were then either sheared for 30 minutes or exposed to static conditions. qPCR was performed. Replicates N = 3. *$p < 0.05$, ***$p < 0.005$. **B.** Western blot demonstrating knockdown of SPTAN1 in THP-1 cells. **C.** Western blot showing knockdown of RAF1 in THP-1 cells. **D.** Western demonstrating decreased STIM1 protein in STIM1-KD THP-1 cells. **E.** Western showing decreased ORAI1 protein in ORAI1-KD THP-1 cells. **F.** Images of the negative controls for the PLA experiments demonstrating SPTAN1 colocalization with KRAS and RAF1.

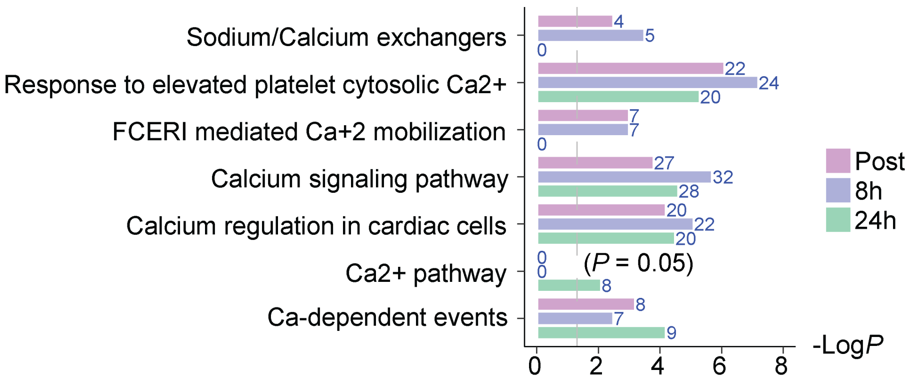


**Figure S7.** Calcium-responsive enrichment pathways for DEGs in snRNA-seq.

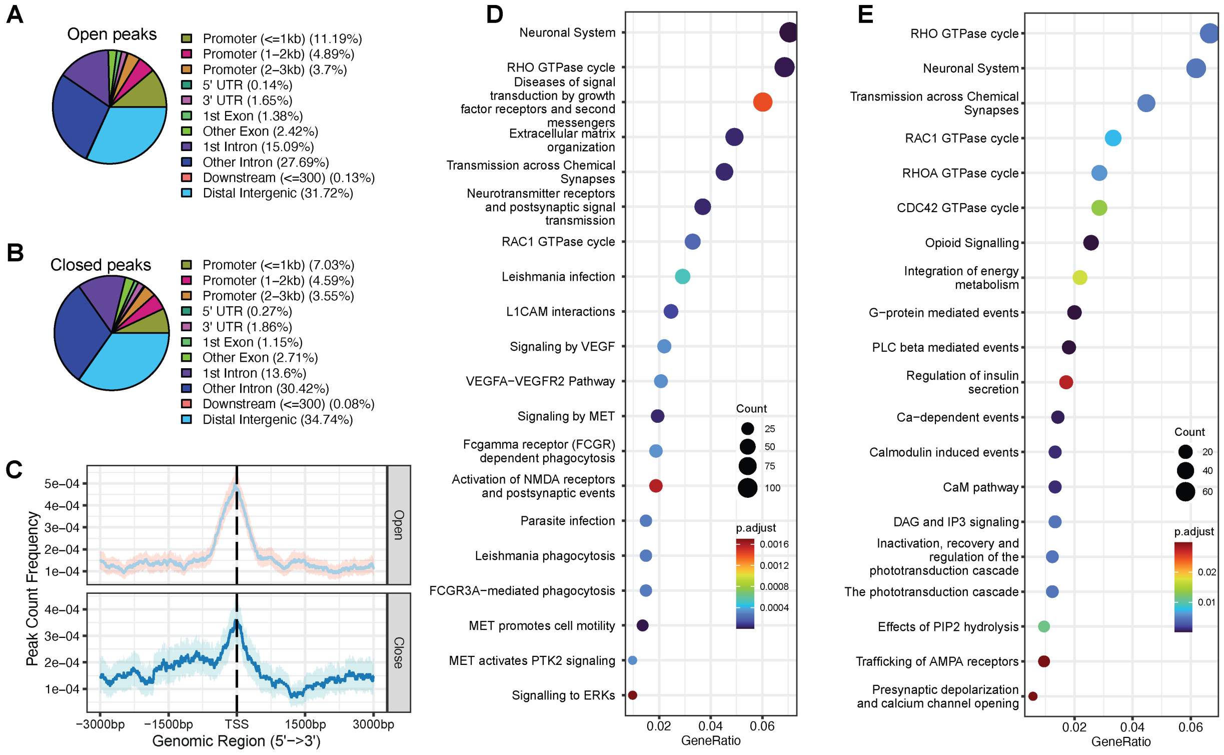


**Figure S8. Structure and functional annotation of differential peaks detected using bulk ATAC-seq in sheared THP-1 cells as compared to static condition. A.** The structure annotation of open peaks in sheared THP-1 cells ($p < 0.05$). **B.** The structure annotation of closed peaks in sheared THP-1 cells ($p < 0.05$). **C.** Average profile of open and closed ATAC peaks binding to transcription starting site (TSS) region with the genomic range from -3000 bp to +3000 bp. **D.** Functional enrichment of open peaks associated gene sets ($p < 0.01$). **F.** Functional enrichment of closed peaks associated gene sets ($p < 0.01$).

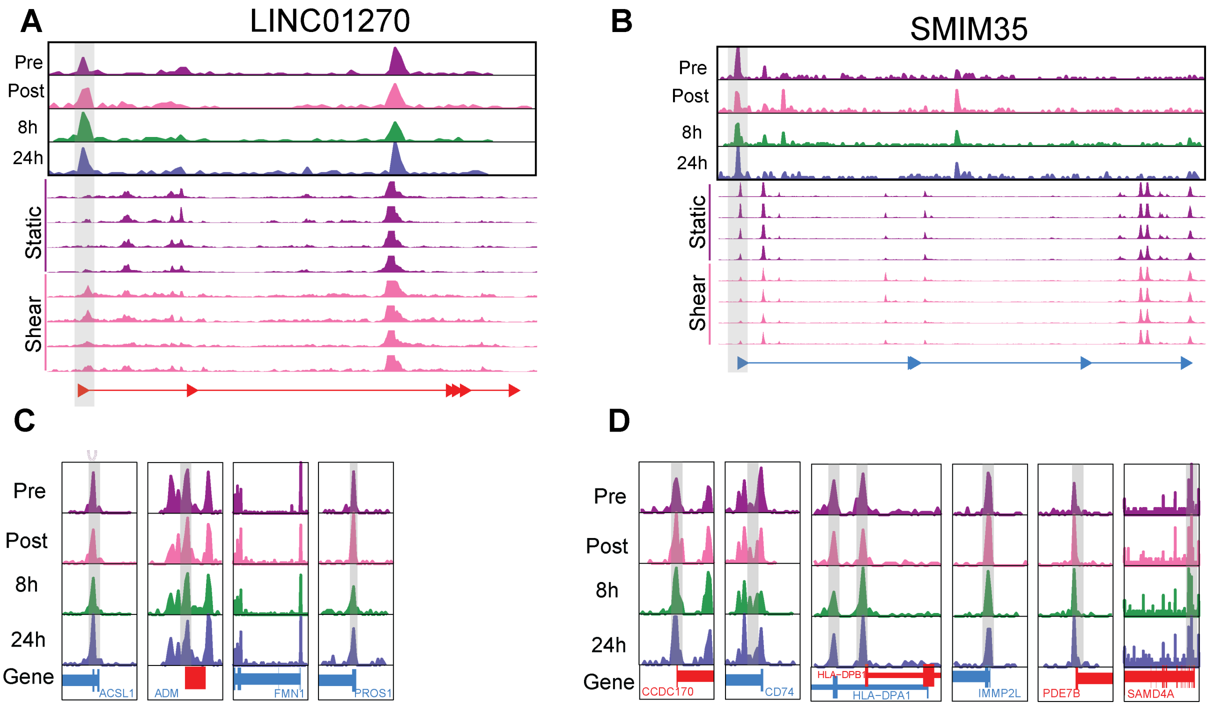


**Figure S9. Important candidate genes with open or closed peaks induced by shear stress at promoter region in classical monocytes of patients and THP-1 cells. A.** Promoter of LINC01270 gene was opened in both classical monocytes of patients and THP-1 cells after shear stress. **B.** Promoter of SMIM35 gene was closed in both classical monocytes of patients and THP-1 cells after shear stress. **C-D.** Four genes were opened, and seven genes were closed at promoter regions in classical monocytes of patients after shear stress.

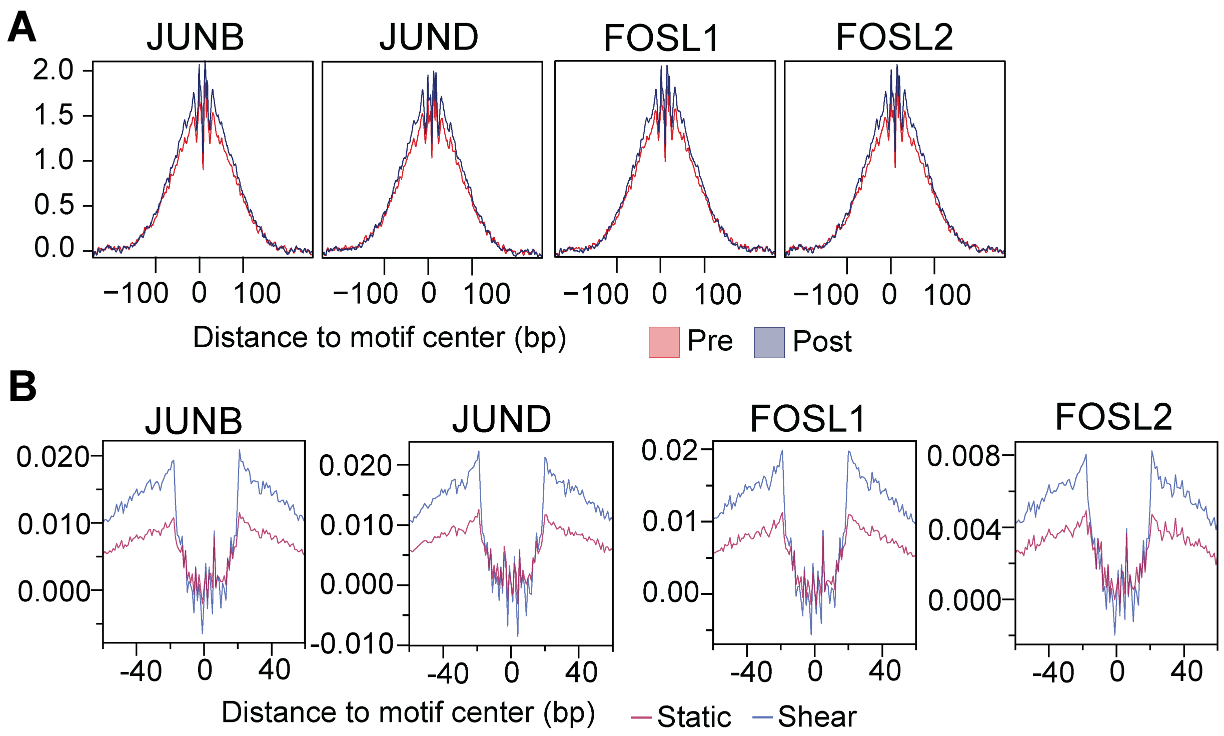


**Figure S10. Transcription factor (TF) footprinting results of JUN and FOS family members between Pre (Static) and Post (Shear). A.** Footprinting signals of JUN and FOS family members increased at Post compared to Pre stage in patients. **B.** Footprinting signals of JUN and FOS family members increased after shear stress compared to static in THP-1 cells.

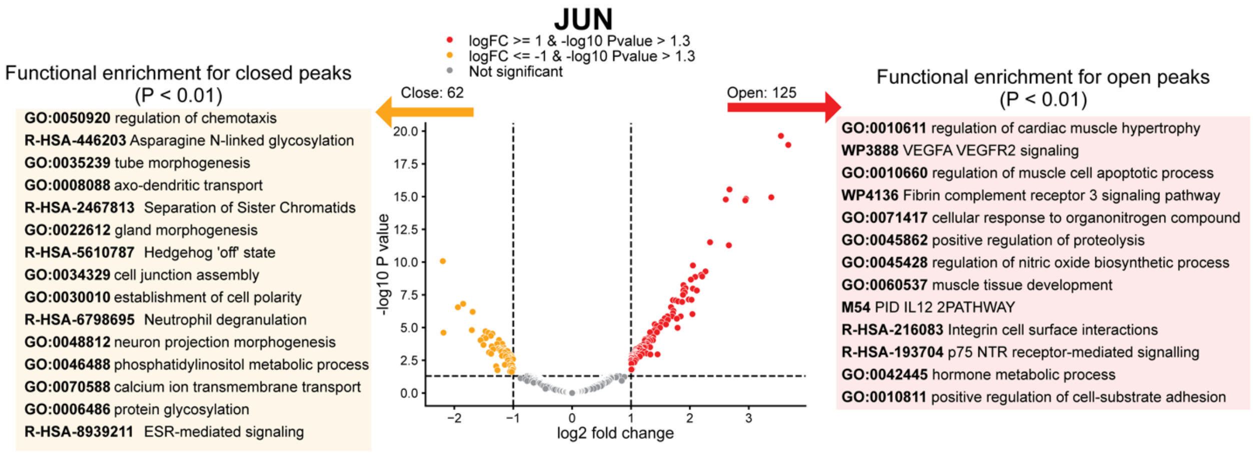


**Figure S11.** Volcano plot visualizing differential peaks detected using CUT&RUN in THP-1 cells treated with transcription factor (TF) JUN antibody. Functional enrichment of open and closed peaks was attached at left and right corner.
